# Oscillatory activity in rostral middle frontal gyrus and subthalamic nucleus encode proactive inhibition in cortico-subcortical motor control network

**DOI:** 10.64898/2026.05.26.728036

**Authors:** Van Tuan Kiet Duong, Seyyed Bahram Borgheai, Enrico Opri, Faical Isbaine, Nicole C. Swann, Nicholas Au Yong, Svjetlana Miocinovic

## Abstract

The current model of the action inhibition network includes the prefrontal cortex and the subthalamic nucleus (STN) connected via the prefrontal hyperdirect pathway. Proactive inhibition refers to preparatory mechanisms that facilitate action inhibition (i.e. enables a person to act with restraint), while reactive inhibition is a sudden stopping triggered by an external stimulus. Most research has focused on the reactive paradigm, with more limited investigation of proactive inhibition. We studied electrophysiologic activity in multiple cortical and STN regions in 17 patients with Parkinson’s disease using high-resolution intracranial electrodes. Subjects performed a Go/NoGo task and a simpler Go task. Proactive inhibition was assessed by contrasting Go trials in the context of two tasks. In the rostral middle frontal gyrus, we found increased beta oscillations during movement preparation in Go trials of the Go/NoGo task compared to the Go task. A similar but weaker, preparatory beta modulation was observed in dorsal STN, while central STN was associated with significant modulation in theta power prior to movement. We interpret this activity as a reflection of the role of these regions in proactively restraining anticipated responses. Conversely, inferior frontal gyrus and ventral STN were primarily engaged during rapid post-cue action control. Specifically, withholding of action after the NoGo signal was accompanied by increased theta activity in these regions. Beta modulation within the STN mirrored those of sensorimotor cortex during successful inhibition and movement execution. In both regions, beta activity decreased during movement and was higher when movement was withheld. We conclude that communication within the hypothesized motor control network is frequency dependent, with key nodes promoting specific functions.

## Introduction

The hypothesized network subserving action inhibition typically includes the (right) prefrontal cortex and the subthalamic nucleus (STN), connected via the prefrontal hyperdirect pathway (Aron, 2011; Haynes and Haber, 2013; Chen et al., 2020). This model is supported by a number of functional and structural studies. For instance, an fMRI study in healthy subjects identified the most prominent structures activated during stopping as the right inferior frontal cortex and the STN, with stopping time inversely correlated with activation levels (Aron and Poldrack, 2006). Similarly, patients with frontal cortex or basal ganglia lesions exhibited significantly longer stopping times compared to healthy controls (Aron et al., 2003; Rieger et al., 2003).

One prevailing theory is that the stopping process begins with the cortex (perhaps the inferior frontal cortex specifically) sending an emergency inhibition signal to the STN, which then suppresses basal ganglia output to prevent movement execution (Aron et al., 2016). Electrophysiological recordings have further demonstrated increased oscillatory activity in the beta band (13–30 Hz) in both the inferior frontal cortex and the STN during stopping tasks, suggesting that components of the action inhibition circuit communicate via the beta band, with the degree of beta power increase linked to inhibitory capacity (Swann et al., 2009; Ray et al., 2012; Aron et al., 2016).

Inhibitory control consists of two components: proactive and reactive inhibition (Aron, 2011). Proactive inhibition is goal-driven, associated with situations where there is knowledge that stopping may be necessary, and it helps control or restrain actions before they begin or to respond more cautiously. Proactive inhibition is particularly relevant for behavioral regulation, as it involves pre-planning based on environmental cues to enhance the likelihood of success. Reactive inhibition, on the other hand, relies on external stimuli, is triggered by a signal indicating that stopping is necessary, and works to halt ongoing actions. Patients with Parkinson’s disease (PD) have impairment in both reactive and proactive inhibitory control, and treatments such as dopaminergic medications or deep brain stimulation (DBS) can further impact these functions and worsen behavioral impulsivity (Jahanshahi, 2013; Deligani et al., 2025). Most electrophysiological and behavioral research of action inhibition has focused primarily on the reactive paradigm, with limited understanding of proactive inhibitory control. Additionally, most prior studies of cortical function have relied on non-invasive recordings such as EEG, which offers limited spatial resolution (Swann et al., 2011; Deligani et al., 2025). Consequently, the specific anatomical substrates in the prefrontal cortex underlying proactive inhibition have not been fully characterized (Swann et al., 2013).

The inferior frontal gyrus (IFG) has been implicated in reactive stopping, potentially triggering stopping via fronto-basal ganglia circuits (Aron et al., 2003; Swann et al., 2009; Aron, 2011), whereas the middle frontal gyrus (MFG), part of the dorsal-lateral prefrontal cortex, is responsible for broader cognitive processes like executive monitoring, working memory, holding a goal in mind, and potentially, proactive control (Chikazoe et al., 2009; Swann et al., 2013; Smith et al., 2019; Khan et al., 2024). The STN is thought to be involved in cognitive control and motor suppression through various cortical and basal ganglia connections (Mink, 1996; Aron et al., 2007; Zavala et al., 2018). Specifically, the ventral regions receiving prefrontal inputs contribute to cognitive control and response inhibition (Haynes and Haber, 2013; Chen et al., 2020), while the dorsal portions project to subsequent nodes in the indirect pathway regulating movement initiation and suppression (Parent and Hazrati, 1995). A better understanding of electrophysiologic changes in the specific regions of the cortex and STN could help clinicians to more optimally apply DBS therapy and specifically avoid inducing impulsivity or even treat executive dysfunction by entraining desired patterns (Ricciardi et al., 2025).

The purpose of this study was to investigate cortical and subcortical electrophysiology during movement execution and proactive inhibition using high resolution intracranial recordings. Defining the organization of the cortex and basal ganglia and characterizing their activity is critical for advancing models of goal-directed cognition. We studied patients with PD undergoing awake DBS surgery and determined electrophysiologic activity in multiple cortical and STN regions of interest during proactive inhibition. Subjects performed two tasks: a Go task and a Go/NoGo task. The Go/NoGo task is a well-established behavioral paradigm to assess a person’s capacity to withhold a prepotent response (Gomez et al., 2007), while the Go task, which lacks NoGo trials, evaluates movement behavior with minimal inhibition. We hypothesized that engaging proactive inhibition is characterized by increased beta oscillations in the IFG, MFG and the STN serving to inhibit action execution in the appropriate context of a Go/NoGo task.

## Methods

### 1. Subject Selection

We recruited patients with idiopathic PD undergoing clinically-indicated STN DBS surgery at Emory University Hospital. All participants provided written informed consent according to the approved Institutional Review Board protocol of Emory University. Rating scale scores, Movement Disorders Society-Unified Parkinson’s Disease Rating Scale Part III (MDS-UPDRS III) and Montreal Cognitive Assessment (MoCA), were obtained during clinical preoperative assessment, approximately 3 months prior to surgery.

### 2. Experimental protocol

#### 2.1 Surgical procedure and recording contact localization

DBS leads (Boston Scientific DB2202 or Medtronic B33005) were implanted either unilaterally or bilaterally into the STN using standard surgical procedure with microelectrode recording guidance (Starr, 2002). To record cortical activity for research purposes, two subdural ECoG strips (28 or 6 contacts; Ad-Tech, Oak Creek, WI) were inserted through the burr hole used for DBS implantation and rested on the cortical surface over one hemisphere (typically right) before clinical microelectrode recordings were started (Panov et al., 2017; Miocinovic et al., 2018; Chen et al., 2020) (Table 1). The 28-contact strip had two rows of fourteen 1.2-mm diameter contacts spaced 4 mm apart. The 6-contact strip had one row of six 5-mm diameter contacts spaced 10 mm apart. One of the strips was targeted to the primary motor cortex (M1) hand knob area while the other strip was targeted to pars triangularis of the IFG using fluoroscopic guidance. In 3 patients ECoG strips were not used due to logistical reasons; additionally, in one patient M1 strip could not be inserted due to high tissue resistance, and in one patient the prefrontal strip was accidentally flipped over and unusable for recordings.

**Table 1.** Clinical characteristics and experimental conditions.

| Patient code | Age (years) | Gender | Disease duration (years) | MDS-UPDRS part3 meds off score | MoCA score | Recording hemisphere | Number of contacts over sensorimotor cortex | Number of contacts over prefrontal cortex |
| --- | --- | --- | --- | --- | --- | --- | --- | --- |
| 1 | 40s | M | 15 | 47 | 30 | R | 28 | 6 |
| 2 | 60s | M | 15 | 43 | 29 | L | 6 | 6 |
| 3 | 50s | M | 8 | 31 | 30 | R | 6 | 28 |
| 4 | 40s | F | 12 | 45 | 26 | R | 28 | 6 |
| 5 | 60s | M | 15 | 34 | 27 | R | NA | NA |
| 6 | 60s | M | 3 | 37 | 23 | R | NA | NA |
| 7 | 70s | M | 18 | 65 | 24 | R | 6 | 28 |
| 8 | 60s | M | 9 | 44 | 27 | R | 6 | 28 |
| 9 | 60s | M | 8 | 32 | 26 | R | NA | 28 |
| 10 | 60s | M | 12 | 33 | 26 | R | NA | NA |
| 11 | 60s | F | 13 | 57 | 22 | R* | 6 | 28 |
| 12 | 60s | F | 6 | 52 | 23 | R | 6 | 28 |
| 13 | 60s | M | 13 | 25 | 29 | R | 28 | 6 |
| 14 | 40s | M | 9 | 38 | 26 | R | 6 | 28 |
| 15 | 40s | M | 5 | 71 | 27 | R | 6 | 28 |
| 16 | 60s | M | 11 | 47 | 24 | R* | 6 | 28 |
| 17 | 50s | M | 8 | 49 | 30 | R | 6 | 28 |
| Average ± STD | 59.2 ± 8.3 | 14 M \ 3 F | 10.6 ± 4.1 | 44.1 ± 12.3 | 26.4 ± 2.6 | 16 R \ 1L | 3 28c \ 10 6c | 10 28c \ 4 6c |

MDS-UPDRS = Movement Disorders Society-Unified Parkinson's Disease Rating Scale; MoCA = Montreal Cognitive Assessment; STN = subthalamic nucleus; STD = standard deviation; R = right; L = left, M = male; F = female; c = contact; NA = not applicable. \*STN signals recorded in the left hemisphere.

ECoG contacts were localized using intraoperative CT images co-registered with a preoperative MRI. For each patient, a three-dimensional mesh model of the brain surface was generated from preoperative T1 MRI using FreeSurfer software(Fischl et al., 2002). Based on the Desikan-Killiany atlas (Desikan et al., 2006), the ECoG contacts were classified as overlying the IFG (pars orbitalis, pars triangularis, pars opercularis), MFG (rostral middle frontal gyrus), premotor (caudal superior frontal and caudal middle frontal gyrus), M1 (precentral gyrus), or S1 (postcentral gyrus). Contacts outside of these regions were not used in the analysis. In one patient, the prefrontal strip was misplaced, overlaying the border between pars opercularis and lateral precentral gyrus, and these contacts were also excluded from analysis. The distance from the ECoG strip to the anatomical midline was measured using the built-in measurement tool in 3D Slicer. A line was drawn from the midpoint of the strip, defined as the midpoint between contacts 3 and 4 for the 6-contact strip, and between contacts 7 and 21 for the 28-contact strip, to the midline in the corresponding MRI scan.

Then, for group visualization purposes, individual patient’s T1 MRI sequences were warped into a normative space known as the Montreal Neurological Institute (MNI) standard brain (Fonov et al., 2009) using SPM12 software accessed through the FieldTrip toolbox (Oostenveld et al., 2011). Individual ECoG contact coordinates were transferred to the MNI space with the normalization parameters using custom MATLAB code and visualized on the reconstructed brain surface using 3D Slicer software (Fig. 1A).

**Figure 1.**
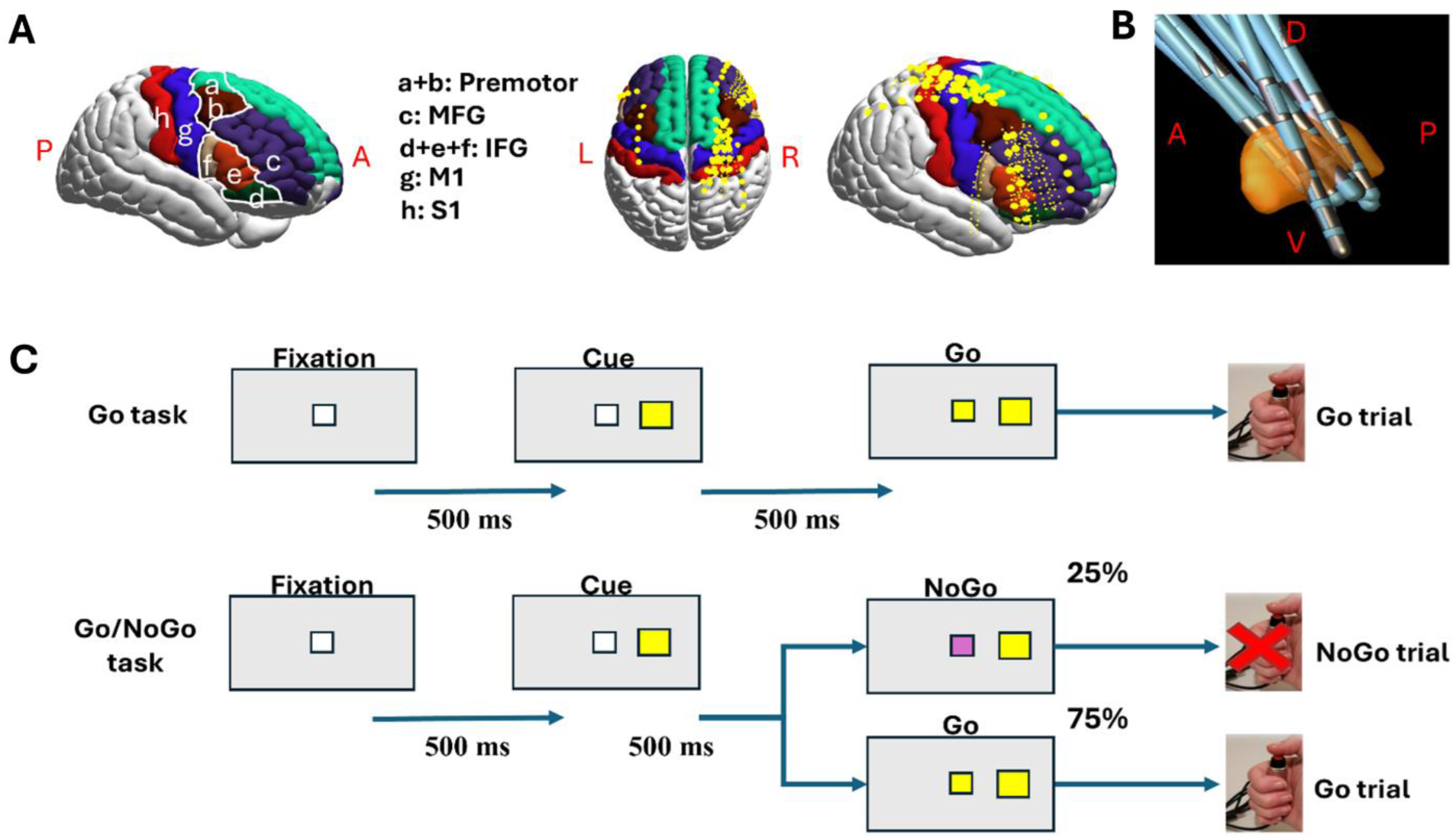
Locations of recording cortical and STN electrodes in a normalized brain and depiction of behavioral tasks. A) MNI brain and normalized position of recording ECoG strips for all patients. (Left) Cortical gyri of interest are color-coded according to Desikan–Killiany atlas in FreeSurfer; a: caudal superior frontal, b: caudal middle frontal, c: rostral middle frontal, d: inferior frontal pars orbitalis, e: inferior frontal pars triangularis, f: inferior frontal pars opercularis, g: precentral, h: postcentral. For spectral analysis, the inferior frontal gyrus (IFG) region was composed of pars orbitalis, pars triangularis and pars opercularis; the middle frontal gyrus (MFG) region included rostral middle frontal gyrus; the premotor region included caudal superior frontal and caudal middle frontal gyri; primary motor region (M1) included the precentral gyrus and primary sensory region (S1) included postcentral gyrus. (Center) Top view of normalized brain with recording ECoG electrodes. (Right) Right hemisphere view of normalized brain with recording ECoG electrodes. B) Anterior view of normalized STN with DBS leads. The bipolar STN channels were grouped as dorsal (D), central and ventral (V). A=anterior; P=posterior; L=left; R=right. C) Behavioral tasks. In the Go task (top), each trial starts with a fixation square, followed 500 ms later by a colored square (Cue signal) on either left or right of the fixation square (corresponding to the hand which will be required to respond). After another 500 ms, the fixation square turns the same color as the cue square requiring a response from the corresponding hand (Go signal). In the Go/NoGo task (bottom), the protocol is similar except that on 25% random trials, the fixation square turns a different color from the cue square, requiring withholding the response (NoGo signal).

DBS electrode contacts were localized on a postoperative CT scan in native (patient-specific) space with CranialSuite software 6.3.1 (Neurotargeting) and referenced to mid-commissural coordinates (Table S1). For group visualization purposes, leads were also localized in normative MNI space using Lead-DBS v3.1 based on postoperative CT and Distal atlas (Horn and Kuhn, 2015; Neudorfer et al., 2023) (Fig. 1B).

#### 2.2 Intraoperative recordings and behavioral tasks

ECoG and LFP were recorded using the Neuro Omega system (Alpha Omega, Nazareth, Israel) at 22 kHz sampling rate with a built-in 0.075-3500 Hz bandpass filter. An ipsilateral scalp needle electrode was used as the reference while contralateral scalp needle electrode served as the ground. Patients were awake (at least 60 minutes after propofol was discontinued) and were approximately 12 hours off dopaminergic medications.

After DBS leads were implanted, patients performed two behavioral tasks, a Go task and a Go/NoGo task (Fig. 1C). The tasks were displayed on a 15.5-inch computer screen approximately 30-40 cm away from patient’s face and angled for optimal viewing while lying semi-recumbent. Patients had a handheld push button switch in each hand and responded by pressing down with their thumbs (Fig. 1C; except one patient used a mouse and pressed right and left buttons with the index and middle fingers of his dominant hand). Button presses were registered through a custom-built Arduino board and sent to both the task computer (via USB port) and Neuro Omega (via BNC cables to the two digital auxiliary channels). Neuro Omega auxiliary signals were sampled at 44 kHz (digitized simultaneously with electrophysiologic signals) and were used to define the response times. Button presses were recorded as HIGH-to-LOW digital state changes. If no button was pressed, there was no recorded state change. To synchronize task events with neural signals, a photodiode, which recorded light changes that corresponded to task events, was affixed to the bottom corner of the screen and recorded with the Neuro Omega system (through the analog auxiliary channel).

Tasks were administered using custom scripts in MATLAB (Mathworks, Natick, MA) (Isoda and Hikosaka, 2007) and Psychtoolbox (Kleiner et al., 2007). Each trial started with the appearance of a fixation signal (a central white 1.8×1.8 cm square). This was followed 500 ms later by a Cue signal (yellow or pink 2.7×2.7 cm square presented to the right or left of the central square) which indicated either a right or left button press would be required. Another 500 ms later, a Go signal was presented which was indicated by the central square changing to the same color as the flanking square. In the Go/NoGo task, on 25% trials, presented randomly, a NoGo signal was indicated by the central square changing to the color that did not match the color of the Cue square (pink or yellow). Patients had 1500 ms to make (or withhold) a response for each trial (Fig 1C). For the Go task, one block of 45 trials was administered after an initial 12-trial practice block. For the Go/NoGo task, 120 trials were administered which were divided into four blocks of 30 trials after a 12-trial practice block (except one patient who performed 90 trials of which 33% were NoGo trials). The tasks were always administered in the same order with the Go task first. The entire protocol took approximately 10 minutes to complete. One patient was not administered the Go task, and one patient declined to perform Go/NoGo task due to fatigue. For two patients, the photodiode sensor came loose during the Go/NoGo task and timings for a portion of trials could not be determined so they were classified as faulty and not analyzed (62 and 12 trials respectively). Additionally, in several patients, a few trials (1-5) lasted longer than the expected 2500 ms due to technical failure and were also classified as faulty and excluded (Table S2).

### 3. Analysis

#### 3.1 Behavioral Analysis

The following behavioral metrics were calculated for each task: median response time (time from Go signal to button press), number of premature error trials (responding after Cue and before, or up to 100 ms after, the Go signal), number of commission error trials (responding on NoGo trials), number of wrong button error trials (responding with the wrong hand) and number of omission error trials (not responding on Go trials). Task accuracy was calculated as the percent of correct trials. Impulsivity on the Go/NoGo task was quantified as the percent of commission errors (number of commission error trials divided by the number of NoGo trials).

#### 3.2 Signal Processing

Data were processed using custom scripts in MATLAB. Electrophysiological signals were band-passed from 2 to 200Hz and notch filtered at 60 Hz using a 3rd order Butterworth filter. Noisy contacts, based on visual inspection, were excluded (1 channel each in 3 patients). ECoG contacts were bipolar montaged by subtracting signals from two directly adjacent contacts (for 6-contact strip) or from two nearby contacts where one contact was skipped (for 28-contact strip). Skipping a contact on the 28-contact strip was performed to make the inter-contact distance similar for the two types of strips (skipped contacts were also bipolar referenced in an interleaved fashion). For DBS lead contacts, segmented contacts on the same level were first averaged and then a bipolar montage was applied to the adjacent contact along the lead. The signals were then epoched into trials using timing derived from the photodiode deflections. The faulty and behaviorally incorrect trials were excluded from analysis. The epoched signals from correct trials were time-locked so that time zero was defined as the Go/NoGo signal.

Time-frequency spectrograms were generated using Morlet wavelets convolution (Cohen, 2019) centered on 200 frequencies from 2 to 200 Hz, with the number of cycles linearly increasing from 3 to 15 such that higher frequencies were associated with more cycles. Individual trial spectrograms were baseline corrected by dividing each data point by the median power during the 300 ms period after the Fixation signal but before the Cue signal (for each frequency separately). Given the difference in number of trials between the Go task and the Go/NoGo task as well as differences in Go and NoGo trial numbers within the Go/NoGo task, we randomly selected the same number of Go trials for each comparative analysis. The spectrogram for each channel was generated by calculating the median of all chosen single-trial spectrograms (median was used to minimize influence of outlier trials). The final spectrogram for each anatomical region was calculated by averaging all spectrograms for that region from all patients. Spectrograms were compared using subtraction and permutation testing with cluster correction as detailed in Statistical analysis section below.

We additionally calculated average power in predefined time-frequency windows relevant for motor control: theta (4-7 Hz), beta (13-30 Hz) and high gamma (100-150 Hz) bands during movement planning period (500 ms before Go/NoGo signal) or movement initiation/inhibition period (100-500 ms after Go/NoGo signal).

#### 3.3 Statistical Analysis

Kolmogorov-Smirnov test was used to determine the normality of the clinical, behavioral and electrophysiological data distribution. Since all data were not normally distributed, non-parametric paired (Wilcoxon signed rank) and unpaired (Wilcoxon rank sum) statistical tests were used for statistical comparisons. Spearman correlation was used for correlations between behavioral and clinical data. A p-value of less than 0.05 was considered statistically significant. Bonferroni correction for multiple comparisons was used when evaluating average time-frequency power in 8 brain regions of interest (significant p-value 0.05/8 = 0.0063).

Non-parametric pixel-wise permutation test and cluster correction were used to compare ECoG and LFP power changes in the spectrograms during different tasks and types of trials (Cohen, 2014). The null hypothesis was that there was no significant difference in power of each pixel between the two conditions (i.e. type of trial). First, the spectrograms in each condition were averaged and subtracted to yield the pixel population that can be used to generate a distribution of differences between the two conditions. The spectrograms of the two conditions were subtracted from one another then multiplied by randomly assigned 1 or −1 (generated based on Rademacher distribution) and then averaged to generate a surrogate difference spectrogram (paired comparison). Multiplication by 1 or −1 is equivalent to swapping labels between the two conditions. The process was then repeated 1000 times to yield a null distribution of difference spectrograms. We called the null distribution spectrograms d_n and our original observed difference spectrogram d_o. To compare each pixel in d_o against the null hypothesis, we computed the z-value of each pixel as: z = (d_o-mean(d_n)) /std(d_n). Any pixel with z-score above or below ±1.96, corresponding to the uncorrected two-tailed p-value of 0.05, were flagged as potentially significant. All remaining pixels were set to zero to generate the thresholded z-map. To account for multiple comparisons, cluster correction was used (Cohen, 2014). A cluster was defined as a collection of pixels that were above the significance thresholds and adjacent to one another. Cluster correction retained only the cluster in d_o whose size (number of pixels) that was larger than 95th percentile of identified clusters in d_n. These qualified clusters were considered significant and are outlined with a black line in figures.

## Results

### 1. Subject Characteristics

A total of 19 patients participated in the study. Data from two participants were excluded due to excessive errors during tasks (>50% incorrect trials), leaving 17 patients in the final analysis (14 men, 3 women; all right-handed; Table 1). The average age was 59.2 ± 8.3 years, the average PD duration was 10.6 ± 4.1 years, the average preoperative MDS-UPDRS III score off-medication was 44.1 ± 12.3, and the average MoCA score was 26.4 ± 2.6. Cortical recordings were obtained from 14 patients (13 right, 1 left hemisphere; Table 1, Table S1). The number of bipolar channels used for spectral group analysis was 26 for IFG, 48 for MFG, 11 for premotor, 17 for M1, 13 for S1, 17 for dorsal STN, 17 for central STN and 17 for ventral STN. STN signals were obtained from the same hemisphere as cortical recordings, except in two patients where the opposite hemisphere was used due to technical issues (Table 1).

### 2. Behavioral Results

Patients performed both tasks with high level of accuracy (92±8% for the Go task and 93±5% for the Go/NoGo task; Table S2). They responded significantly faster during the Go task (median response time 370.1 ± 123.7 ms) compared to the Go/NoGo task (median response time 444.1 ± 104.5 ms) (p=0.042; Fig. 2A; Table S1). Longer response times were expected since proactive inhibition was engaged during the Go/NoGo task. The Go/NoGo commission error percentage did not correlate with the response time prolongation (difference between Go/NoGo and Go response times), suggesting that impulsivity on the Go/NoGo task was not a purely motor phenomenon. In other words, since the error rate was not directly related to the degree of proactive motoric slowing, other cognitive processes were likely involved to generate impulsive action (Fig. 2B). In fact, motor disease severity was significantly correlated with response time in the Go task but not in the Go/NoGo task (Fig. 2C). Conversely, MoCA score, a measure of global cognitive function, was negatively correlated with response time in the more complex Go/NoGo task but not in the simple Go task (Fig. 2D). There was no correlation between the MoCA score and response time prolongation or MoCA score and accuracy for either task (not shown).

**Figure 2.**
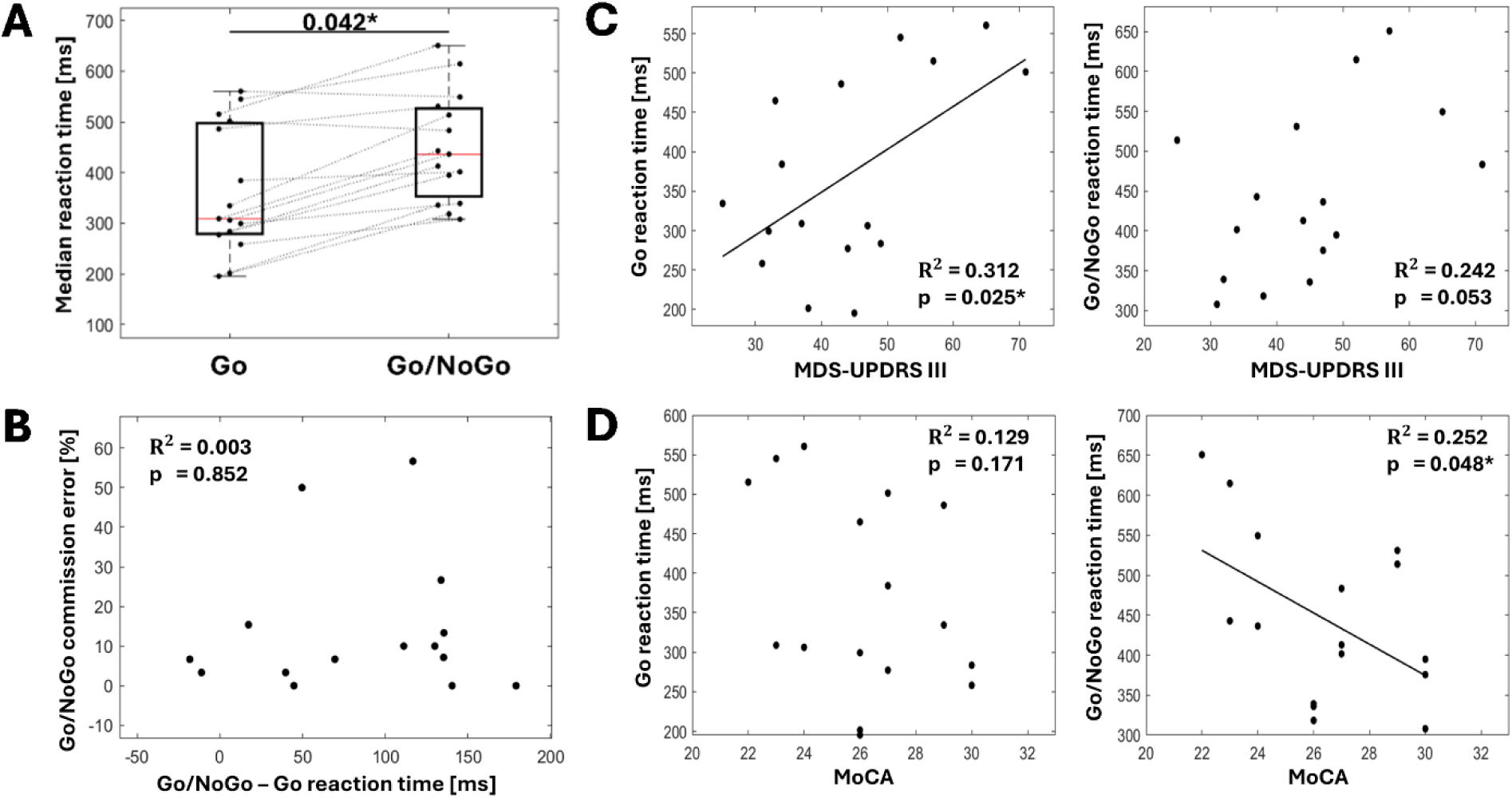
Behavioral results. A) Response time was significantly longer for Go trials from the Go/NoGo task compared to the Go task. B) Difference between the Go/NoGo and Go task median reaction time (response time prolongation) was not correlated with commission error percentage (impulsivity). C) The MDS-UPDRS III off medication score was positively correlated with Go task response times (left) but not with Go/NoGo task response times (right). D) The MoCA score was negatively correlated with Go/NoGo task response times (right) but not with Go task response times (left).

### 3. Time-Frequency Results

#### 3.1 Spectral power changes between the Go trials of the Go/NoGo task and Go task

We compared spectral changes during movement preparation and successful execution between the two tasks to determine the effect of proactive inhibition in each anatomical region of interest. Since proactive inhibition was engaged during the Go/NoGo task but not for the Go task, spectral activity during the Go trials of the Go task was subtracted from the activity during Go trials of Go/NoGo task which allowed us to evaluate the effect of proactive control (Fig. 3).

**Figure 3.**
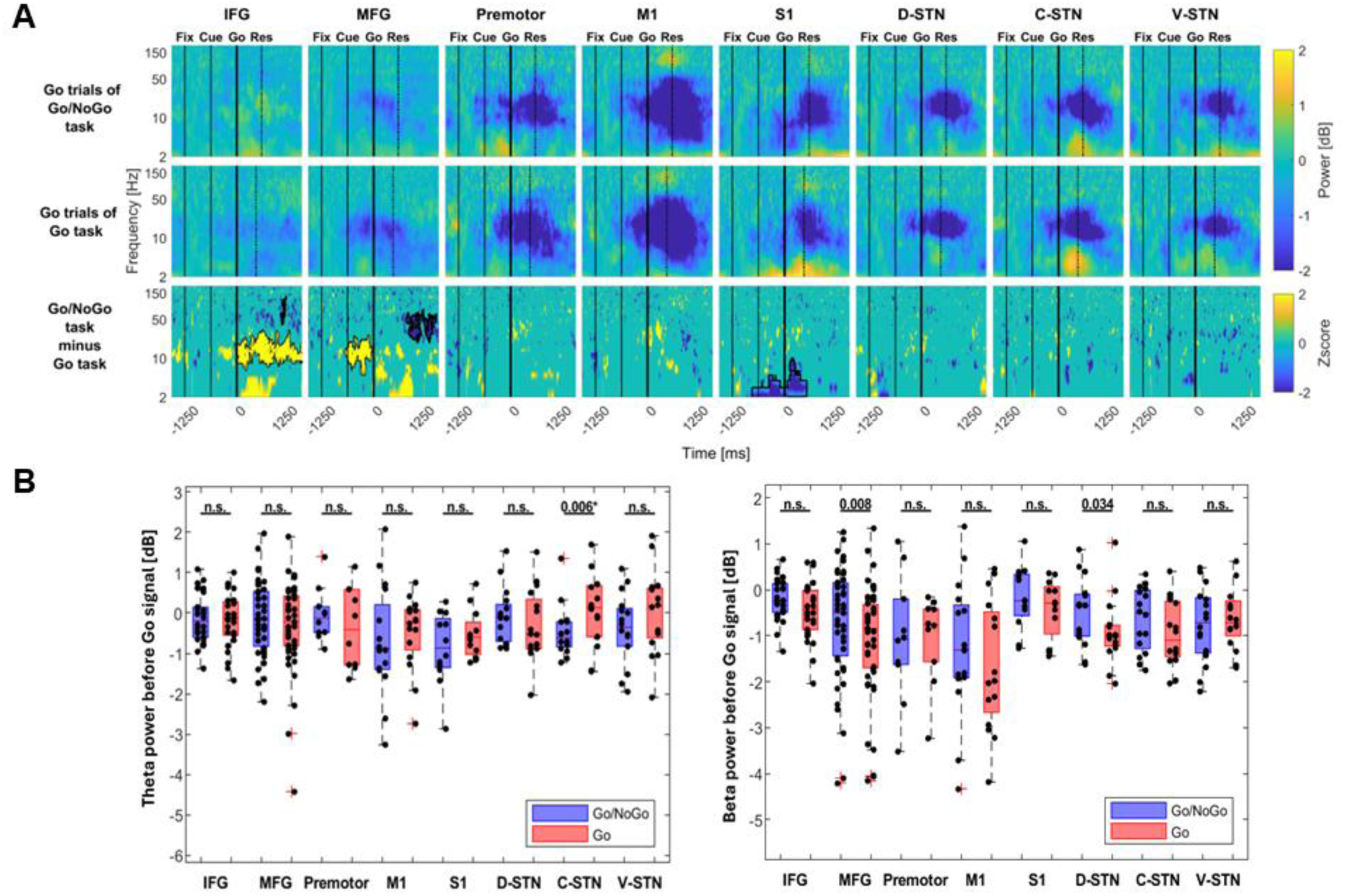
Proactive inhibition assessment: ECoG and LFP spectral power comparison between Go trials in Go/NoGo and Go tasks. A) Averaged time-frequency spectrograms of channels overlaying different cortical regions (inferior frontal gyrus [IFG], rostral middle frontal gyrus [MFG], premotor, primary motor [M1], primary sensory [S1]) and within the STN regions (dorsal [D-STN], central [C-STN], ventral [V-STN]) are shown for the Go trials of the Go/NoGo task (top row), Go task (middle row), and their z-scored difference (bottom row). The non-zero threshold for the z-score map is set at |z|>1.96, corresponding to an uncorrected p-value<0.05 (meaning all non-zero pixels are significant but before cluster correction). Solid black borderline indicates statistically significant difference after cluster correction. Time zero is the Go signal (thick black vertical line) with Fixation and Cue signal times indicated by black solid vertical line and the median response time indicated by black dotted vertical line (variable between subjects). B) Boxplots of average spectral power for each channel during movement preparation period (500 ms between Cue and Go signals) for theta (4-7 Hz) band (left) and beta (13-30 Hz) band. On each box, the central mark indicates the median, and the bottom and top edges of the box indicate the 25th and 75th percentiles; the whiskers extend to the most extreme data points not considered outliers, and the outliers are plotted using the ‘+’ symbol. Asterisk indicates significant p-value after multiple comparison correction (0.05/8 regions). N.s. = not significant.

In the IFG, despite both types of trials resulting in movement, there was a significant increase in IFG beta power immediately *after* the Go signal as well as around and after the response time in the Go/NoGo task (Fig. 3A). A small period of increased alpha/beta power *before* the Go signal (movement preparation period) was not significant after cluster correction. After the response, gamma power (∼50-150Hz cluster) was transiently higher during the Go task compared to Go/NoGo task.

In the MFG, beta power decreased less before the Go signal (movement preparation period) in the Go/NoGo task compared to Go task, resulting in significantly higher power after subtraction (Fig. 3A). This difference also had a trend toward significance when considering average beta power before the Go signal (p= 0.008; Fig. 3B). This pattern during the movement preparation period may be indicative of proactive inhibition processes engaged during the Go/NoGo task that are absent during the Go task. After the response, beta and low gamma power (∼30-100 Hz cluster) were significantly higher during the Go task compared to Go/NoGo task.

In the premotor and M1 cortex, there was no significant difference in spectral power between Go/NoGo and Go tasks (Fig. 3A). In the S1 cortex, increased power in low frequency (2-8 Hz) cluster was present during movement planning and around the response time in the Go task (Fig. 3A). This may be due to sensory feedback from holding the thumbs firmly against the buttons, which was likely more pronounced during the Go task when movement execution was always expected. This finding was not present when trials were aligned to the response time confirming that it was likely related to anticipatory preparation before the Go signal (Fig. S1).

In the STN, during movement preparation period (before the Go signal), average theta power was significantly lower during the Go/NoGo task (p=0.006; Fig. 3B). There was also a trend toward higher average beta power in the dorsal STN during Go/NoGo task (p=0.034; Fig. 3B). On visual inspection, theta power was increased around the response time (most prominently in the central STN), which was comparable for the two tasks. This may reflect general sensorimotor processing of the response across tasks.

#### 3.2 Spectral power changes between the Go and NoGo trials of the Go/NoGo task

We investigated the activity in each anatomical region during movement execution and inhibition by comparing correct Go and correct NoGo trials of the Go/NoGo task (Fig. 4).

**Figure 4.**
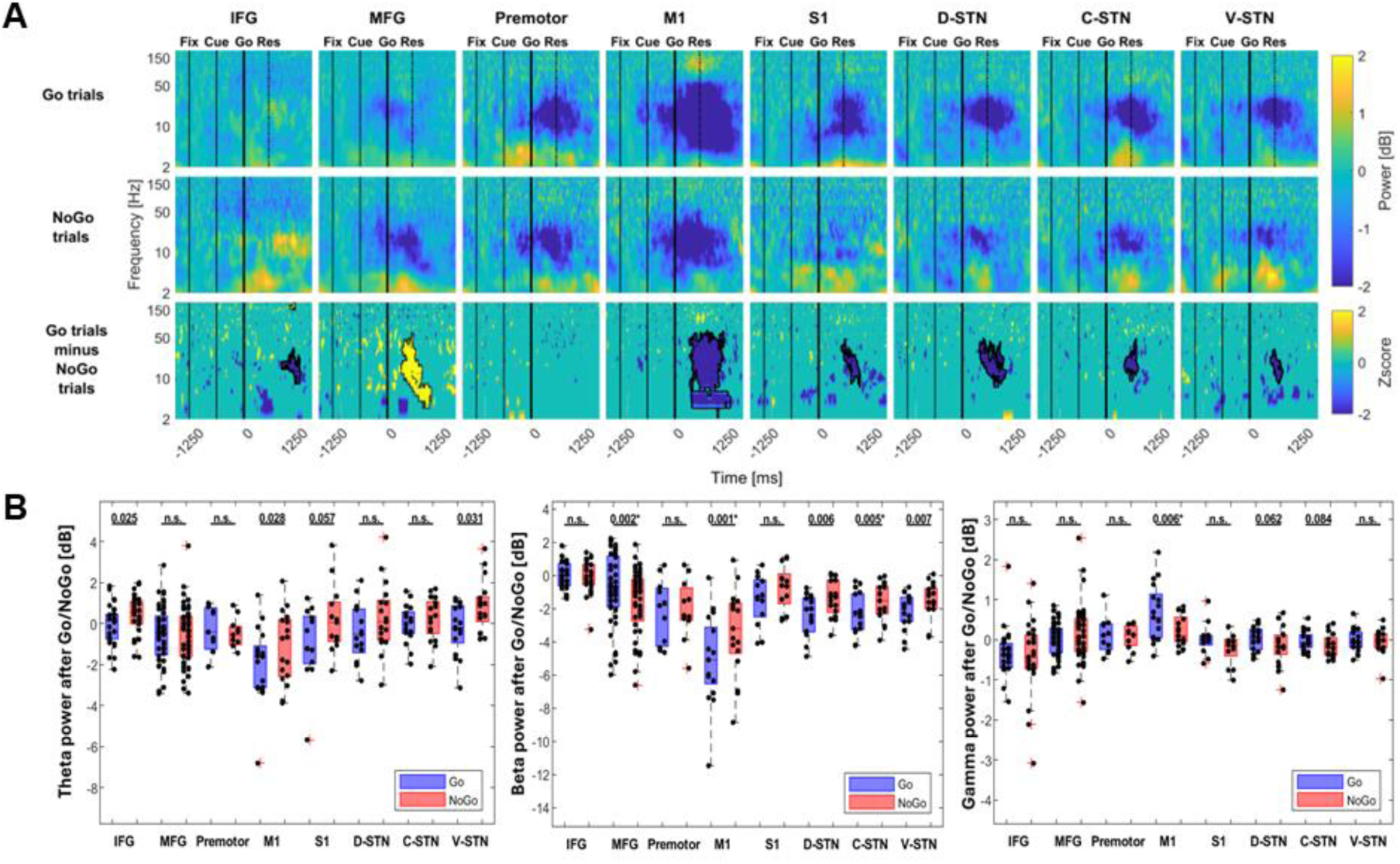
Reactive inhibition assessment: ECoG and LFP spectral power comparison between Go trials and NoGo trials in Go/NoGo task. A) Averaged time-frequency spectrograms of channels overlaying different cortical regions (inferior frontal gyrus [IFG], rostral middle frontal gyrus [MFG], premotor, primary motor [M1], primary sensory [S1]) and within the STN regions (dorsal [D-STN], central [C-STN], ventral [V-STN]) are shown for the Go trials (top row), NoGo trials (middle row), and their z-scored difference (bottom row). The non-zero threshold for the z-score map is set at |z|>1.96, corresponding to an uncorrected p-value<0.05 (meaning all non-zero pixels are significant but before cluster correction). Solid black borderline indicates statistically significant difference after cluster correction. Time zero is the Go (top) or NoGo (middle) signal (thick black vertical line) with Fixation and Cue signal times indicated by black solid line and the median response time indicated by black dotted line (variable between subjects). B) Boxplots of average spectral power for each channel during movement initiation or inhibition period (100-500 ms after the Go/NoGo signal) for theta (4-7 Hz) band (left), beta (13-30 Hz) band (middle) and high gamma (100-150 Hz) band (right). On each box, the central mark indicates the median, and the bottom and top edges of the box indicate the 25th and 75th percentiles; the whiskers extend to the most extreme data points not considered outliers, and the outliers are plotted using the ‘+’ symbol. Asterisk indicates significant p-value after multiple comparison correction (0.05/8 regions). N.s. = not significant.

In the IFG, there was an increase in beta power during later period of the NoGo trials (Fig. 4A). This delayed activity may reflect feedback for correctly inhibiting a response. Immediately after the NoGo signal, theta power was increased compared to the same time period after the Go signal. This spectral cluster was not significant after correction, but the difference trended toward significance when comparing average power (p=0.025; Fig. 4B).

In the MFG, there was a larger alpha/beta power decrease during NoGo compared to Go trials, which was opposite to the pattern observed in the sensorimotor brain regions (M1, S1, STN) where a larger beta power decrease was observed during the Go trials. This unexpected decrease in beta band was also significant when comparing average power (p=0.002; Fig. 4B).

In the M1 and S1, there was a more profound decrease in beta power during movement execution compared to NoGo trials, while no differences were observed in the premotor cortex. In M1, average gamma power increased around the movement execution time compared to the NoGo trials (p=0.006; Fig. 4B) while theta power after NoGo trended toward an increase (p=0.028; Fig. 4B).

In the STN, beta power decrease was more pronounced when there was movement execution compared to the NoGo trials in all STN subregions (Fig. 4A; p=0.006, p=0.005, p=0.007; Fig. 4B). There was a trend toward higher theta power after the NoGo signal compared to the Go signal in ventral STN (p=0.031; Fig. 4B).

## Discussion

We studied electrophysiological changes in multiple cortical and subcortical structures during movement execution and inhibition in PD patients at high spatial and temporal resolution. We hypothesized that proactive inhibitory control would be characterized by increased beta oscillations in the IFG, MFG and the STN. Behaviorally, we observed that subjects exhibited longer response times in the cognitively more demanding Go/NoGo task requiring proactive inhibition compared to the simple Go task. Different brain regions had distinct oscillatory patterns when proactive inhibitory processes were engaged and when movement was executed or inhibited. Our findings suggest that in prefrontal cortical regions, increase MFG beta band activity is predominantly related to proactively restraining anticipated responses during the preparatory period (i.e. holding goals in mind) thereby confirming our hypothesis, while IFG was mostly oriented towards rapid action control after the command signal which was manifest through increased theta activity. In the STN, there was some evidence of increased beta power in dorsal STN in maintaining proactive inhibitory control, similar to what was observed in MFG, while central STN had decreased theta activity. The sensorimotor cortex, on the other hand, was primarily involved in movement initiation and execution, which was manifested through beta band desynchronization. Beta modulation within the STN was found to mirror that of sensorimotor cortex during both successful inhibition as well as movement execution. Theta activity increased after the Go or NoGo signal in multiple regions, suggesting a role in rapid action control, but this may also reflect signal detection or increases in attention, depending on context. Overall, we find that communication within the hypothesized motor control network is frequency dependent, with theta and beta activity in MFG and dorsal/central STN reflecting engagement of proactive inhibitory control while IFG and ventral STN modulate theta activity during reactive inhibition.

### Rostral middle frontal gyrus beta synchronization: a hallmark of proactive inhibition

Electrophysiological and tracing studies have identified a fronto–basal ganglia network involving the IFG and dorsolateral prefrontal cortex (DLPFC) that connect to ventral STN through the prefrontal hyperdirect pathway to regulate inhibitory control (Swann et al., 2009; Haynes and Haber, 2013; Chen et al., 2020; Mosher et al., 2021). Most high-resolution electrophysiological recordings have been focused on reactive inhibition through the implementation of the stop signal task, leaving unanswered questions on the electrophysiological mechanisms required for more cognitively demanding proactive inhibition. Both reactive and proactive inhibitory processes are thought to rely on similar neural substrates (Swick et al., 2011; Aron et al., 2016), communicating through beta (Swann et al., 2009; Chen et al., 2020; Deligani et al., 2025) and low-frequency oscillations (Zavala et al., 2018; Khan et al., 2024).

In our study, we observed that prefrontal beta power was higher during the Go trials of Go/NoGo task compared to Go task in both IFG and MFG. However, the time course of the increase was different between the two. In MFG beta was primarily modulated during the preparatory period (between Cue and Go) while in IFG beta was modulated during the execution period (after the Go signal). This points to functional differences between the two anatomical structures. The DLPFC, which the MFG is a part of, is known to participate in diverse cognitive functions, including high-level executive tasks like maintaining goal-related working memory and inhibiting prepotent action (Chikazoe et al., 2009; Degutis et al., 2024). On the other hand, IFG, especially the right hemisphere IFG, has been implicated in reactive inhibition, rapidly initiating a stopping command, a process accompanied by elevated beta power following an external stop cue (Swann et al., 2009; Swann et al., 2012; Aron et al., 2016). Additionally, IFG has elevated theta activity in the Go/NoGo task after both the Go and NoGo signals. Based on prior work which have implicated prefrontal theta oscillations in cognitive control this increase could reflect additional cognitive processing engaged to discriminate between Go and NoGo trials (Helfrich and Knight, 2016; Pagnotta et al., 2024). In the STN, proactive inhibition was associated primarily with *decreased* theta activity in the central region (at the interface between motor and non-motor STN subdivisions) and possibly higher beta activity in the dorsal motor region (mirroring the MFG findings). Increased theta activity has been associated with inhibitory control in the cortex, but in the STN, increased theta power is associated with response execution (Zavala et al., 2018). So, our findings suggest that anticipation of a definite response (Go task) is similarly reflected in higher theta power in the STN. Overall, the evidence suggests proactive inhibitory control relies on the modulation of beta oscillatory activity in the MFG, with additional contribution from the STN in the beta and theta bands (Fig. 5).

**Figure 5.**
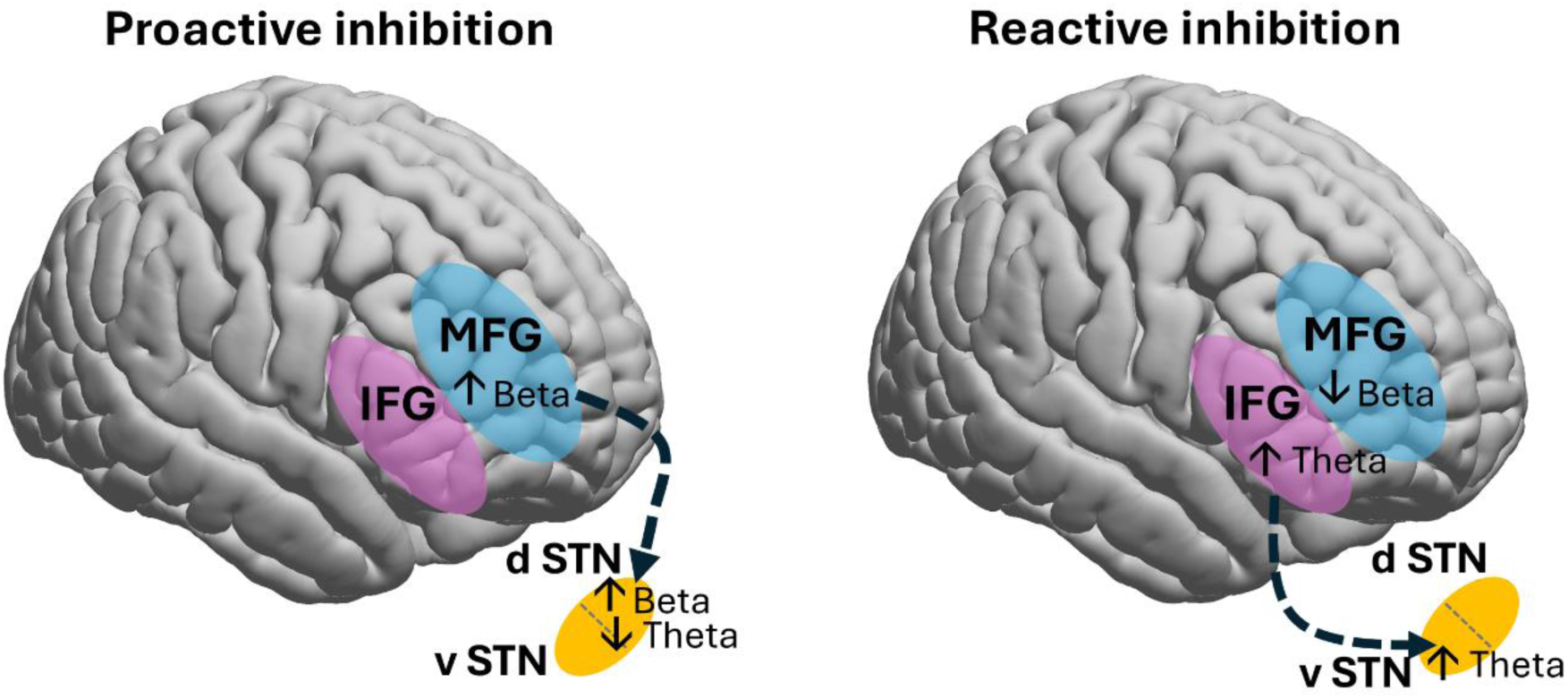
Schematic of the hypothesized inhibitory motor control network. The network subserving action inhibition includes the (right) prefrontal cortex and the subthalamic nucleus (STN), connected via the prefrontal hyperdirect pathway. Distinct regions exhibit specific oscillatory dynamics depending on whether proactive inhibitory control is engaged. The rostral middle frontal gyrus (MFG) supports proactive response restraint with increased beta activity during movement preparation, but decreased beta power during response withholding. The inferior frontal gyrus (IFG) is involved in reactive inhibition via increased theta activity. Dorsal (d) and central STN regions are engaged in proactive control through increased beta and decreased theta oscillations. Ventral (v) STN increases theta oscillations during response withholding.

### Prefrontal and subthalamic oscillatory dynamics in response inhibition

While the Go/NoGo task primarily probed proactive inhibition by requiring participants to maintain preparatory inhibition in anticipation of infrequent NoGo trials, reactive inhibition was also present when the NoGo signal appeared (Aron, 2011). From previous studies, beta oscillations are known to increase in the IFG (Swann et al., 2009; Swann et al., 2012; Chen et al., 2020) and the STN (Kuhn et al., 2004; Wessel et al., 2016; Zavala et al., 2018) when individuals cancel or withhold their action following an external cue. Our results also revealed higher beta power for NoGo trials compared to Go trials of Go/NoGo task in both IFG and STN. However, their time course was different. In the IFG, we found that significant beta synchronization happened after the median response time. Unlike the Stop Signal task, our study participants were not moving when the NoGo signal appeared, indicating that the late beta synchronization may therefore indicate that a response was successfully withheld (there were too few incorrect NoGo trials for analysis of unsuccessful inhibition). This difference in timing is also consistent with ‘pause-then-cancel’ model which suggests that there is an initial pause to stop a movement and then a later inhibitory function to retain this behavior (i.e. keep movement withheld) (Schmidt and Berke, 2017; Diesburg and Wessel, 2021). In the STN, there was greater beta desynchronization centered around the median response time in Go trials compared to NoGo trials. MFG, on the other hand, had more alpha and beta desynchronization in NoGo trials compared to Go trials around the median response time, similar to what has been shown previously in DLPFC (Khan et al., 2024). This is consistent with the idea that in prefrontal cortex, alpha and beta oscillations are hypothesized to play a role in working memory control (Schmidt et al., 2019; Pagnotta et al., 2024). The desynchronization of alpha and beta in MFG after NoGo signal could therefore reflect a mechanism of inhibition, allowing access to relevant working memory information, such as the current trial is a NoGo trial and requires continuous withholding of movement, while inhibiting interference from competing information. This is complementary to early (before Go/NoGo signal) MFG beta band activity in the Go trials within the Go/NoGo task which could reflect retention of information and inhibition of competing information from disrupting the working memory goal. These findings suggest that IFG activity was primarily associated with successful post-cue action control, which was distinct from the preparatory action control observed in MFG.

Synchronization in the theta band was observed after the NoGo signal in the IFG and ventral STN which could indicate a potential communication mechanism in cancelling prepotent action. Our results in prefrontal cortex agreed with a previous high-resolution cortical recording study using Go/NoGo task, which show mid-rostral DLPFC low-frequency activity increase when there was a correct inhibitory response (Khan et al., 2024). Low-frequency oscillations in the STN are also known to increase when individuals are making decisions in the presence of inhibition or conflict (Harmony et al., 2009; Zavala et al., 2018). Previous experiments have also revealed low-frequency coherence between STN and various regions of the prefrontal cortex during response inhibition tasks (Zavala et al., 2018; Chen et al., 2020). Together, these findings suggest that cognitive processing and communication between the prefrontal cortex and STN may be encoded in different frequency bands (primarily theta and beta), in the context of successfully withholding a prepotent action (Fig. 5).

### Subthalamic and sensorimotor cortex activity in movement initiation and execution

Ample research has characterized the time course and pattern of neural activity in the sensorimotor cortex during movement execution. Prior to movement, sensorimotor beta power is reduced and remains desynchronized until movement termination. After movement, beta activity bounces back, overshooting the baseline then decreasing back to baseline level (Kilavik et al., 2013; Heinrichs-Graham and Wilson, 2016; Deligani et al., 2025). This pattern is also known to occur in the STN (Kuhn et al., 2004). Additionally, during movement, high gamma activity increases briefly and is localized to the region of the M1 responsible for the specific movement (Crone et al., 1998; Cheyne et al., 2008).

In our study, sensorimotor cortex and STN oscillatory activity during movement is consistent with prior research. The primary sensory, motor cortex, and STN were dominated by beta desynchronization centered around median response time regardless of presence or absence of proactive inhibition. Beta oscillations were desynchronized during NoGo trials as well, but to a lesser degree than when movement was executed. In the STN, the dorsal region exhibited the biggest difference in beta desynchronization between Go and NoGo, which is consistent with the dorsal STN’s role in motor control (Mosher et al., 2021). There was also no gamma synchronization in M1 during NoGo trials, so this finding is specific to the execution of movement. Overall, these results demonstrate that STN and sensorimotor beta oscillation patterns are quite similar when it comes to movement initiation and execution. It is likely that these regions are coordinating their activity through modulation of the beta band.

### Limitations

There are several limitations to our study, largely due to the inherent constraints of intraoperative invasive recordings in human participants. First, the final sample size was relatively small (n = 17), not all patients had recordings available from all three regions of interest (prefrontal cortex, sensorimotor cortex, and STN), and primarily the right hemisphere activity was recorded. Second, the anatomical locations of the ECoG strips and DBS leads were not identical in all patients. The averaging of recordings across slightly different sites to achieve sufficient statistical power could, therefore, introduce spatial effects into our analysis. Third, the task order was fixed (Go task, followed by GoNoGo task) due to operational constraints which may have led to partial confounds of adaptation and fatigue. Finally, all participants were individuals with PD, a population known to exhibit impaired inhibitory control and altered electrophysiological patterns (Gauggel et al., 2004; Meyer et al., 2020; Deligani et al., 2025). As such, the observed electrophysiology may reflect pathophysiology, which limits the generalizability of our findings regarding proactive inhibitory control to healthy populations.

## Conclusion

This study examined high resolution electrophysiological activity across cortical and subcortical regions during movement execution and proactive inhibition in PD patients. Prefrontal regions showed distinct roles; specifically, the MFG supported proactive response restraint during preparation, while the IFG was involved in rapid post-cue action control, primarily via beta band synchronization and transient increases in theta activity. STN was engaged in proactive control through theta and beta oscillations and was active during response withholding. Our results show that distinct brain regions exhibit specific oscillatory dynamics depending on whether proactive inhibitory control is engaged or movements are executed or suppressed. Our findings further elucidate inhibitory motor control network and may help inform more targeted and effective therapeutic application of DBS for PD.

## Data and Code Availability

Available upon request with a formal data sharing agreement.

## Author Contributions

S.M., N.C.S. designed research; S.M., E.O., F.I., N.A.Y., S.B.B. performed research; N.C.S. contributed unpublished reagents/analytic tools; V.T.K.D., S.B.B. analyzed data; V.T.K.D., S.M. wrote the first draft of the paper; N.C.S., E.O., F.I., N.A.Y., S.B.B. edited the paper.

## SUPPLEMENTARY MATERIALS

**Table S1.** Recording contact locations.

| Patient code | Sensorimotor ECoG<br>distance from midline<br>(mm) | Prefrontal ECoG<br>distance from midline<br>(mm) | STN lead bottom<br>contact AC-PC<br>coordinates (mm) |
| --- | --- | --- | --- |
| 1 | 34.2 | 30.63 | 9.6, -4.0, -5.7 |
| 2 | 39.48 | 53.97 | -11.7, -5.1, -6.3 |
| 3 | 24.32 | 50.49 | 11.0, -4.4, -5.1 |
| 4 | 31.69 | 49.85 | 9.6, -3.3, -7.9 |
| 5 | NA | NA | 11.7, -0.6, -3.9 |
| 6 | NA | NA | 9.7, -3.5, -5.4 |
| 7 | 45.33 | 59.34 | 8.0, -3.0, -7.4 |
| 8 | 27.83 | NA | 11.4, -1.7, -4.6 |
| 9 | NA | 57.84 | 10.0, -3.3, -5.1 |
| 10 | NA | NA | 11.7, -1.3, -5.0 |
| 11 | 22.25 | 42.11 | -10.8, -4.1, -4.1 |
| 12 | 32.63 | 49.42 | 9.0, -3.4, -3.8 |
| 13 | 30.81 | 49.65 | 8.1, -4.4, -5.7 |
| 14 | 32.21 | 47.58 | 10.3, -4.9, -5.3 |
| 15 | 29.01 | 51.18 | 12.1, -2.0, -6.5 |
| 16 | 23.62 | 47.16 | -12.0, -0.6, -5.3 |
| 17 | 35.11 | 41.19 | 10.2, -2.5, -5.1 |
| Average ±<br>STD | 31.4 ± 6.4 | 48.5 ± 7.5 | 10.4±1.3, -3.2±1.3, -5.4±1.1 |

ECoG = electrocorticography; STN = subthalamic nucleus; AC =anterior commissure; PC = posterior commissure; STD = standard deviation; NA = not applicable.

**Table S2.**
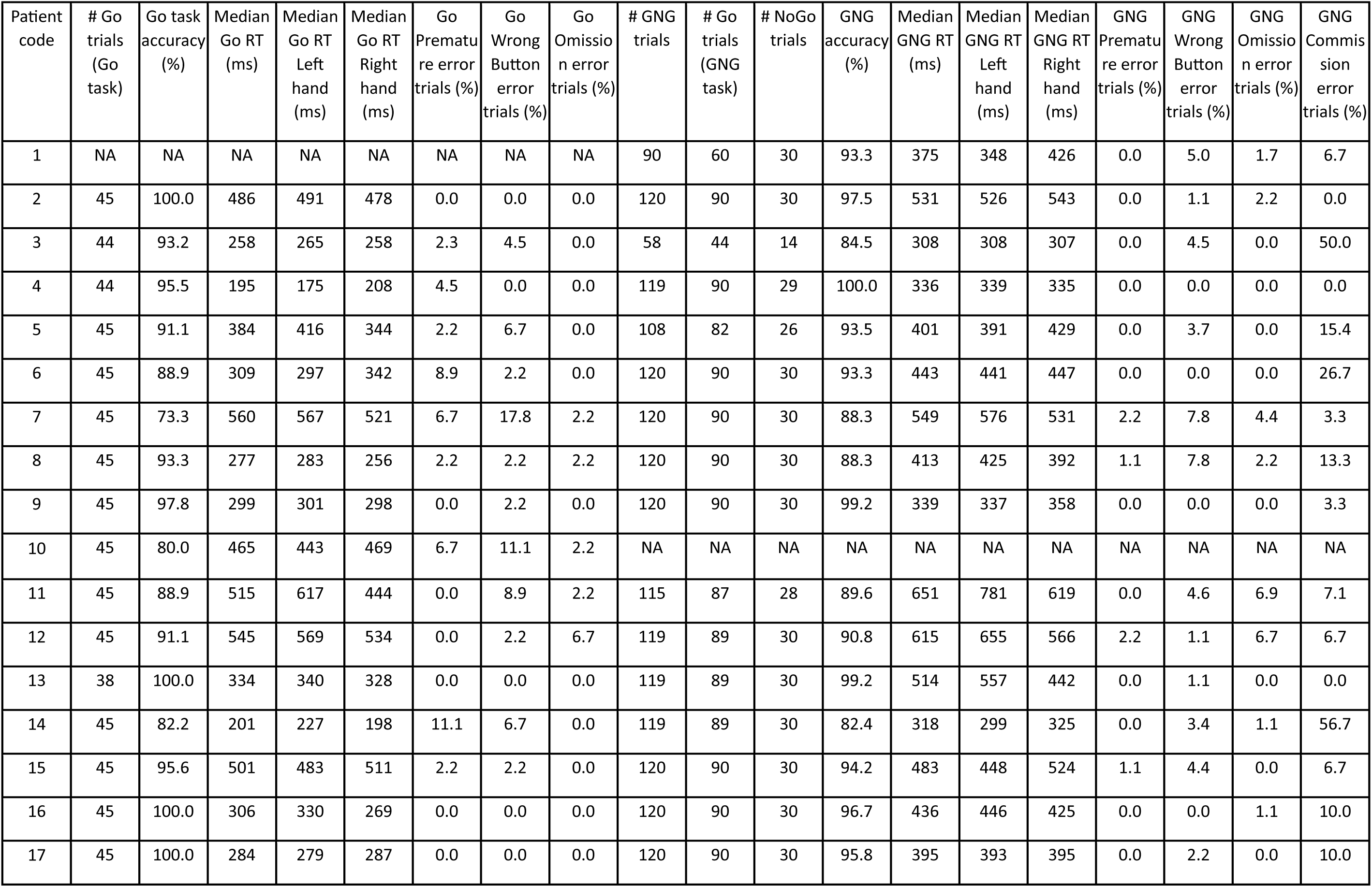
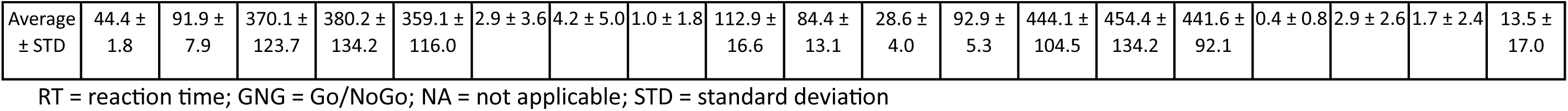
Behavioral performance on Go and Go/NoGo tasks.

**Figure S1.**
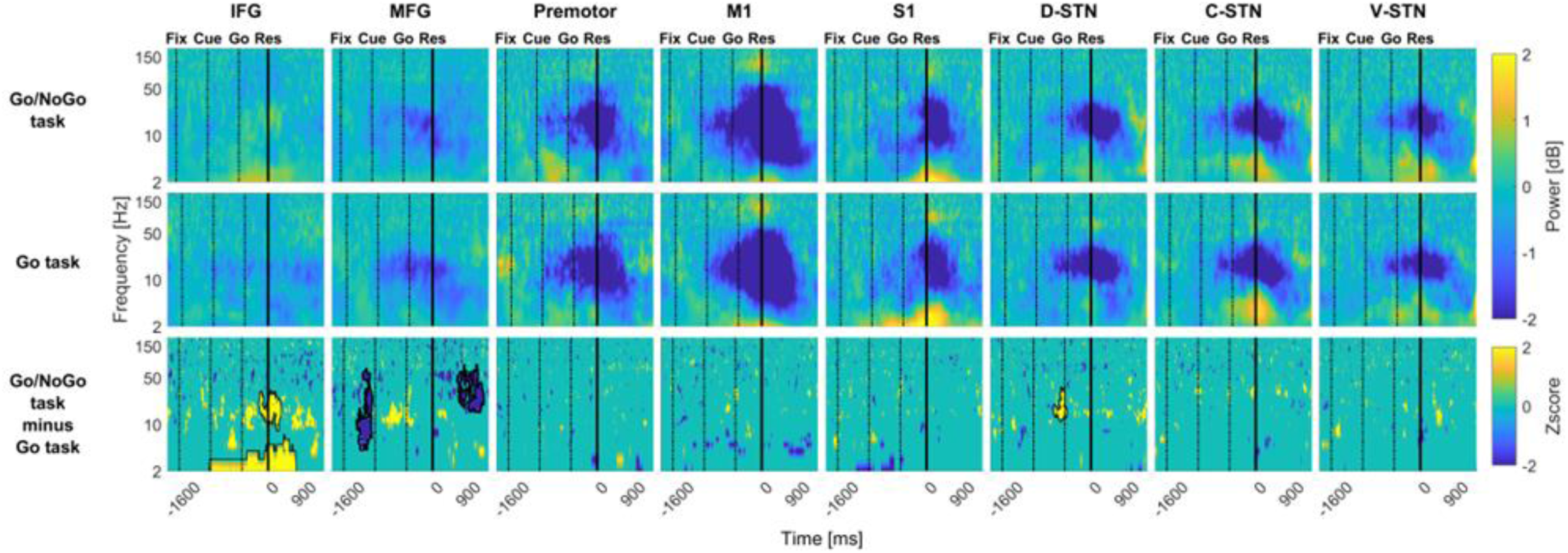
ECoG and LFP spectral power comparison between Go trials in Go/NoGo and Go tasks. Same as Figure 3 but trials aligned to the response. Averaged time-frequency spectrograms of channels overlaying different cortical regions (inferior frontal gyrus [IFG], rostral middle frontal gyrus [MFG], premotor, primary motor [M1], primary sensory [S1]) and within the STN regions (dorsal [D-STN], central [C-STN], ventral [V-STN]) are shown for the Go trials of the Go/NoGo task (top row), Go task (middle row), and their z-scored difference (bottom row). The non-zero threshold for the z-score map is set at |z|>1.96, corresponding to an uncorrected p-value<0.05 (meaning all non-zero pixels are significant but before cluster correction). Solid black borderline indicates statistically significant difference after cluster correction. Time zero is the median response time (thick black vertical line) with Fixation, Cue and Go signal times indicated by black dotted vertical lines (variable between subjects).

